# Effects of Maize Development and Phenology on the Field Infestation Dynamics of the European Corn Borer (Lepidoptera: Crambidae)

**DOI:** 10.1101/2024.01.29.577014

**Authors:** Sacha Revillon, Christine Dillmann, Nathalie Galic, Cyril Bauland, Carine Palaffre, Rosa Ana Malvar, Ana Butron, François Rebaudo, Judith Legrand

## Abstract

Phenological match/mismatch between cultivated plants and their pest could impact pest infestastion dynamics in the field. To explore how such match/mismatch of plant and pest phenologies may interact with plant defense dynamics, we studied the infestation dynamics of maize by one of its main pests in Europe, the European Corn Borer (*Ostrinia nubilalis*; Lepidoptera: Crambidae). A two-year field experiment was carried out on a collection of 23 maize inbred lines contrasted for their earliness. Each inbred line was sown at three different dates in order to expose different developmental stages of maize to natural European corn borer infestation. The effect of the sowing date depended on the inbred line, the pest generation and the year. In 2021, the final pest incidence ranged from 36% to 91% depending on inbred lines and sowing date. In 2022, it ranged from 2% to 77%. This variability in final pest incidence can be related to variations in plant development during plant exposure to pest infestation. However, this relationship was not straightforward. Indeed, the shape and intensity of the relationship depended on the timing of the onset of the pest infestation. When infestation occurred while plants were in a vegetative stage, a non-linear relationship between development and pest incidence was observed with the least and most developed plants being the most infested. When infestation occurred when all plants were in the mature phase, the most developed plants were the least infested. Our results highlight the effect of plant-pest phenological match/mismatch on pest infestation dynamics and underline the importance of taking plant-pest interactions into account to propose relevant control strategies.

## Introduction

The interactions between species are stage dependent and vary throughout their respective life cycles and phenology (Yang and Rudolf, 2010). This is notably the case for herbivorous insects, where host seeking and feeding behaviours are specific to certain life stages (Hagstrum and Subramanyam, 2010). In the European corn borer, *Ostrinia nubilalis* Hübner (Lepidoptera: Crambidae), a pest of maize (*Zea mays* L.), neonates have a pre-feeding dispersal phase while late instars bore into maize stems in order to develop and survive winter (Ross and Ostlie, 1990). The timing of the phenological events in plant and insects is influenced by genetic, abiotic (Ausin et al., 2005) and biotic factors (Hanley and Fegan, 2007; Rasmussen and Yang, 2022). For maize, the timing of events is also determined by agricultural practices, such as sowing dates.

The impact of phenology on species interactions has been studied in the context of climate change, as rising temperatures and the increase in frequency of extreme meteorological events can shift the phenology of species. This shift may differ among species (Parmesan, 2007) and thus could desynchronize interacting species and consequently impact their interactions. When resource availability is limited in time, the demand by the consumer must be synchronized with the provider (Cushing, 1990; Bewick et al., 2016; Ferreira et al., 2023). This is the case in herbivorous interactions, where individuals requiring a resource for their development must be able to feed when this resource is available. For example, an early post-winter resurgence of the herbivorous *Operophtera brumata* Linnaeus (Lepidoptera: Geometridae) prior to the vegetative recovery of its host can lead to the starvation of the neonates and thus a decrease in fitness (Tikkanen and Julkunen-Tiitto, 2003).

Plant-pest interactions are also impacted by the defense strategies developed by plants in response to predation during plant-pest co-evolution. There is a large body of work investigating plant defense mechanisms from a range of perspectives from molecular genetics to physiology and ecosystem ecology, and many hypotheses have been formulated regarding the mechanisms for reducing herbivory and its impact (Stamp, 2003; Gong and Zhang, 2014). Plant defense mechanisms have been classified into three groups (Painter, 1951): (i) tolerance mechanisms that minimize the impact of herbivore damage (ii) antixenosis that reduces the probability of contact between pests and plants, and (iii) antibiosis, i.e. the ability of a plant to reduce or stop the growth and/or development of the pest. Plant defense traits include both physical structures (e.g. spine, leaf toughness) and chemical compounds such as synthesised toxins or volatile organic compounds (Mithöfer and Boland, 2012) that can be constitutive and/or inducible, i.e produced in response to pest infestation. Specifically, in maize, lignin content is a morphological trait whose concentration increase with development and also in response to insect feeding (Meihls et al., 2012; Rodríguez et al., 2012). Defense traits can also be related to dispersal, plant size, or phenology (Boege and Marquis, 2005).

Thus, there is a whole range of defense mechanisms and the resulting level of plant defense can vary. Indeed, plant defense mechanisms may vary during plant development, between individuals due to genetic variation, and with environmental conditions due to phenotypic plasticity (Ballhorn et al., 2011). Hence, genetic variation and temporal variation are two different components of plant defenses that may interact with the match/mismatch of pest and plant phenologies, with contradictory consequences in terms of pest attack dynamics. Plant defenses tend to increase with physiological development until maturity and can later either decrease or be maintained depending on the reproduction strategy (Boege and Marquis, 2005). However, meta-analyses aiming to characterize general patterns of changes in plant defense during development showed that this can vary between species, within species, and with the type of defense (physical or chemical) (Barton and Koricheva, 2010; Quintero et al., 2013, Cao et al. 2019). Temporal changes in physical and chemical defenses may result in non-linear dynamics of defense acquisition. For example, the concentration of DIMBOA (benzoxazinoid), one of the main constitutive chemical defenses in maize leaves, decreases as the plant matures while physical defenses such as lignin and silica concentration (cell wall components) tend to increase over time (Meihls et al., 2012).

The production of these defenses requires resources. However, plants must simultaneously ensure at least three key functions to maximize their fitness: growth, reproduction, and defense, potentially leading to a trade-off between defense and growth (Herms and Mattson, 1992; Züst and Agrawal, 2017). This implies that, within a species, plant defenses could be higher in slow-growing genotypes. This growth-defense trade-off has been the subject of several studies with contrasted results depending on the species considered (Todesco et al., 2010; Stowe and Marquis, 2011; Bode and Kessler, 2012; Santiago et al., 2016). This trade-off could interfere with changes in plant defense during development during development and affect plant-pest interactions.

Due to the interplay between (i) changes in plant defense during development, (ii) the trade-off between growth and defense in the host plant and (iii) the match/mismatch of plant and pest phenologies, the success of the pest on an inbred line can depend on the sowing date and the environmental condition of the season. Within the framework of crop pest control, understanding how the interplay between pest phenology and host plant development in the field could lead to the development of alternative control strategies.

Here, we focus on the interaction between maize crop and the European corn borer (*Ostrinia nubilalis)*. European corn borer is a major arthropod pest of maize in Europe and can infest up to 60% of fields and cause losses of 5% to 30% (Meissle et al., 2009, Kaçar et al., 2023). Studies of the impact of maize earliness and/or phenology on corn borer infestation success have produced contradictory results. A study by Ordas et al. (2013) showed that, within a panel of early and late inbred lines derived from a common ancestral line, early inbred lines suffered less damage from borers,in terms of tunnel length, ear damage. This result can be explained by the more advanced developmental stage and greater tissue maturity of early lines at the time of infestation. By contrast, another experimental study showed that late maturing inbred maize lines displayed better pest resistance (Schulz et al., 1997). This could be explained by a growth-defense trade-off or other genetic correlations between resistance and developmental traits. Other studies explored the role of plant development by shifting the sowing date in order to synchronize or desynchronize the most susceptible stage of crop development and the peak of pest attack, and found that late-sown plants suffered higher yield loss (Malvar et al 2002) and higher ear damage (Blandino et al, 2008). Maize growth stage has been notably identified as a factor involved in the European corn borer oviposition preference. The first generation was reported to prefer early sowings in which the plants are at an advanced vegetative stage, while the second generation of borers, which causes the most damage to plants, prefers physiologically younger plants (Huber et al. 1928, Balachowsky 1966). Chemical compounds in the leaves, such as fructose, whose quantities vary with development, have been correlated with oviposition preference (Derridj et al. 1989).

These previous studies have quantified the impact of pests on plants through harvest observations providing integrated traits of plant resistance or tolerance. However, their contrasting results could be due to variations in the dynamics of pest attacks over the season. Therefore, in order to dissect the trajectories leading to pest infestation severity at the end of the season and to highlight how the interplay between changes in plant defense during development and the phenological match/mismatch between plants and pests impacts pest success, we monitored the infestation dynamics in parallel with plant development. Using a panel of inbred maize lines that differ in their phenology, we investigated how natural field infestation dynamics were driven by plant development, earliness and their interplay with the timing of the onset of the pest infestation.

## Materials and methods

### Plant material

Plant material consisted of a panel of 23 maize inbred lines with contrasting values for several traits including flowering time (Table 1). Thirteen inbred lines, for which leaf secondary metabolites and physiological traits at two developmental stages were previously characterised (Cañas et al., 2017), were selected from a core collection representing European and North American maize genetic diversity (Camus-Kulandaivelu et al., 2006). Three additional inbred lines (Cm484, F271, F66) were chosen for their differences in cell wall digestibility (lignin, hemicellulose or p-coumaric acid content) (El Hage et al., 2018; Zhang et al., 2019) and three other inbred lines were chosen for their tolerance (F618) or susceptibility (Mo17, B37) to the European corn borer (Panouillé et al., 1998; Willmot et al., 2004). Finally, two pairs of sister-inbred lines derived from two ongoing divergent selection experiments (DSEs) for flowering time were included (Durand et al., 2010). Each DSE was initiated from an initial inbred line (F252 or MBS). Cycles of selection for earliness or lateness were carried out from each ancestral inbred line by applying divergent selection. One inbred line derived from the 18^th^ selection cycle for earliness and one derived from the 18^th^ selection cycle for lateness from each initial inbred line were included in the panel (early FP036 and late FT318 from F252, and early MP052 and late MT040 from MBS). The two pairs of DSE-derived lines are therefore genetically close but differ in their days to flowering.

**Table 1.**
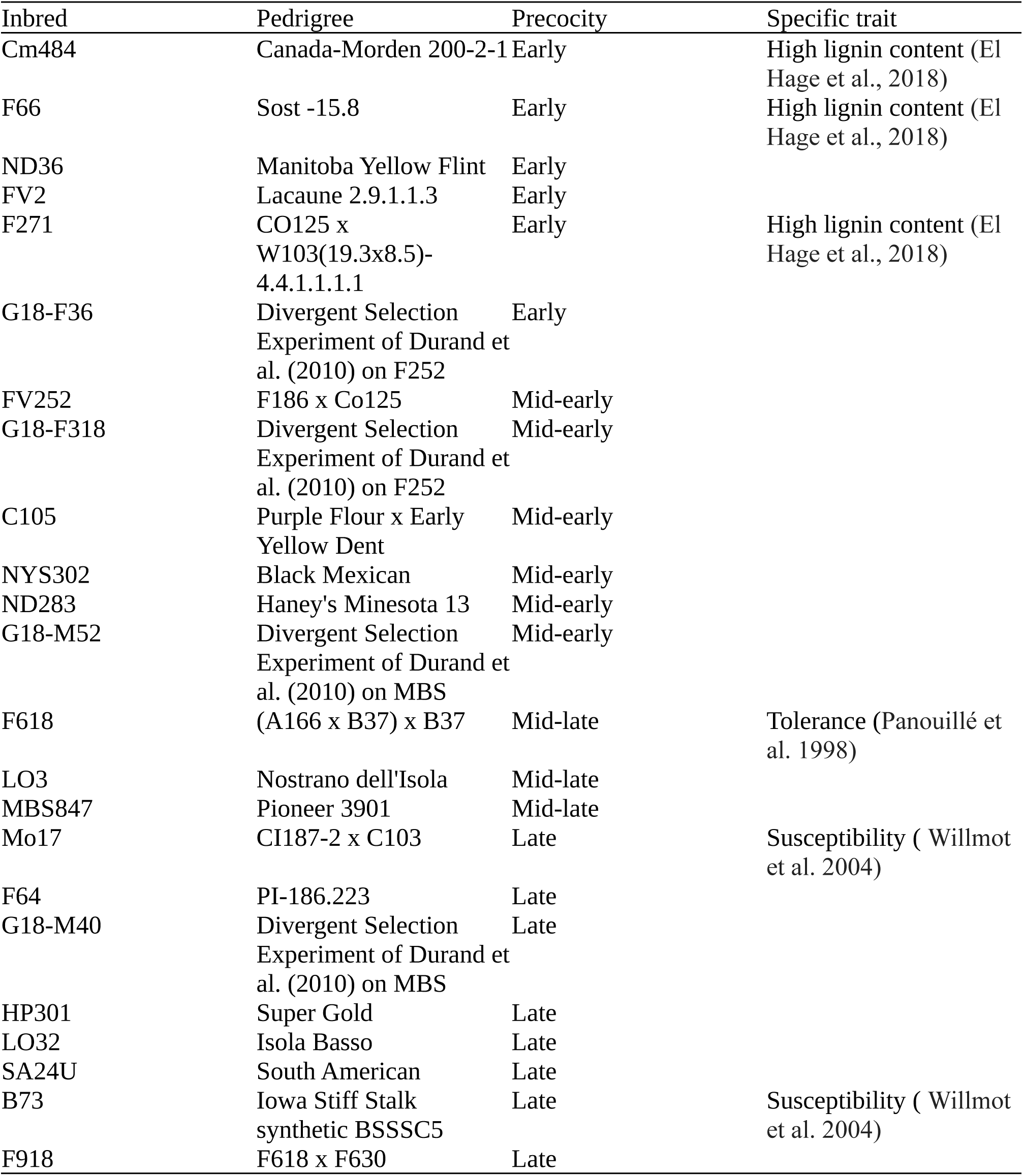
Maize panel description.

### Field experiment

A field experiment was carried out in 2021 and 2022 in Gif-sur-Yvette, France (N 48°717401’, E 2°153018’). In both years, inbred lines were sown at three sowing dates (blocks) (24 April, 12 May and 15 May 2021; 28 April, 11 May and 13 May 2022) in order to expose plants at different developmental stages to the pest. Each block was divided into thirteen sub-blocks (0.80m apart) of sixteen plots (0.80m apart) (Fig. S1). Each inbred line was assigned to a different plot consisting of a single row. Twenty-five seeds were sown in each plot. The first and last plot of each sub-block, as well as the first and last sub-block of each block, were used as borders and sown with MBS847. At each sowing date, each inbred line was sown in six plots (replicates). The six replicates for each inbred line at each sowing date were randomized in the blocks with the condition that the same inbred line must not be sown more than twice in one sub-block. Due to a lack of seeds, inbred lines ND283 and ND36 could not be sown in 2021 and were replaced by G18-M52 from the DSE. Experiments were performed under natural infestation.

### Data collection

#### Plant development and phenology

To assess plant phenology, the dates of female flowering (silking) and male flowering (tasseling) were recorded. The date of silk/tassel emergence corresponded to when 50% of plants in a plot showed silks/tassels.

#### Pest infestation dynamics and pest development

In order to assess the natural infestation dynamics of European corn borer in the field, weekly non-destructive plant monitoring was carried out between mid-June and mid-September. A plant was considered to be infested when it had at least one of the following signs: one or more series of rectilinear holes on the leaves made by the larvae (Fig. S2a), holes and larval frass on the tassels (Fig. S2b), stems (Fig. S2c), and/or ears, and/or a broken stem (Fig. S2d). A plant recorded as being infested on a given date was not longer monitored after that date. In order to determine European corn borer voltinism, two sources of data were used. First, five to ten plants from the MBS847 inbred line sown at the first sowing date were dissected each week and examined for the presence of pupae. Second, data collected in the Ile-de-France region by a national pest surveillance system (Bulletin de Santé du Végétal n°32, 28/09/21 and 20/09/22) were used to assess the number of pest generations and the approximate flight dates.

### Statistical analyses

In order to study the effects of the dynamics of plant development on the dynamics of pest infestation, parameters describing both dynamics were defined for each plot in the field. We describe below how we characterised plant phenology, earliness and pest infestation dynamics. Data from each year of the experiment were analysed separately due to contrasting climatic conditions (cold spells and excessive rainfall in 2021, heat waves and drought in 2022), which led to different developmental dynamics for both plant and pest.

#### Characterising maize phenology

For each plot, phenology was quantified as the flowering date (FD) in days after 1 July. Earliness was quantified by computing the thermal time between sowing and flowering (TTF), i.e a standardised measure of the number of days of development at 20°C required to flower (Parent et al. 2010). Tasseling and silking were characterized separately.

#### Characterising infestation dynamics

To characterise the dynamics of pest infestation on each inbred line for each sowing date, we first estimated the key dynamics parameters at the plot level. Over the two years of the experiment, we observed contrasted infestation dynamics, reflecting the differences in the climatic conditions of 2021 and 2022. In particular, from the survey of MBS plants, we observed a single European corn borer generation in 2021 compared to two generations in 2022. Bivoltinism in 2022 resulted in two periods of infestation (referred to here as the first and second pest generation), compared to univoltism that resulted in a single period of infestation in 2021 that was more spread out over the season than each of the two infestations of 2022. Thus, for a given plot, the European corn borer infestation dynamics were quantified by the incidence at time *t (t* being the calendar date), noted I_t_, and defined as the proportion of infested plants within a plot at time *t*. Incidence data were fitted to a single logistic growth equation for the 2021 data and a double logistic growth equation for the 2022 data (Fernandes et al., 2017) using a non-linear regression:

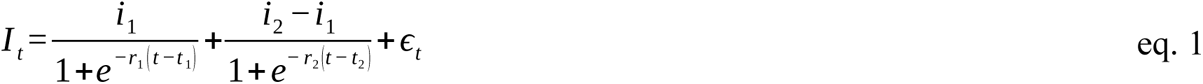

To obtain a single logistic growth equation, the second term of eq. 1 is omitted. Parameter *i_1_* corresponds to the maximum incidence at the end of the first wave of infestation; parameter *i_2_* corresponds to the maximum incidence at the end of the second wave. Thus, the quantity *i_2_ – i_1_* corresponds to the incidence that is exclusively due to the second pest generation. Parameters *r_1_* and *r_2_* are the slope parameters of the first and second waves, respectively. Parameters *t_1_* and *t_2_* are the inflection points of the incidence during the first and second waves, respectively. Each of these points corresponds to the date at which the incidence reaches half of *i_1_*(half of *i_2_*, respectively). *ε* _t_ are Gaussian homoscedastic residuals (Fig. S3). These parameters (*i_1_*, *i_2_ – i_1_*, *r*_1_, *r*_2_, *t*_1_ and *t*_2_) characterize the infestation dynamics and were estimated for each plot. Both single and double logistic growth equations (corresponding to one or two infestation waves, respectively) were fitted for each plot in 2022. The best fit between single and double logistic growth was selected both with the deviance information criterion (DIC) and by visual inspection. For both years, the date of the first attack on the plot *t_0_* was also recorded.

#### Identifying the sources of variation

All traits related to maize phenology and infestation dynamics were fitted to a Gaussian conditional autoregressive (CAR) model that includes inbred line and sowing date factors and accounts for spatial autocorrelation. Model parameters were estimated in a Bayesian framework (Lee, 2013):

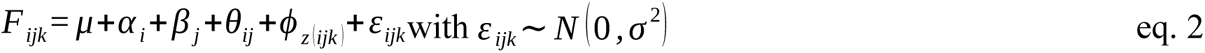

where F_ijk_ is the dependent quantitative variable (either a plant or an infestation-related trait) of the k^th^ plot sown with inbred line *i* at sowing date *j*, μ is the overall mean, *α_i_*is the i^th^ inbred line effect, *β_j_* is the j^th^ sowing date effect, *θ_ij_* is the term of the interaction between *α_i_* and *β_j_*, *ϕ_z(ijk)_* is the spatial effect defined by equation 3 in (Leroux et al., 2000), and ɛ_ijk_ are the residuals. Independent Gaussian priors were defined for *α_i_*, *β_j_,* and *θ_ij_*, and uniform priors for ρ and τ^2^.

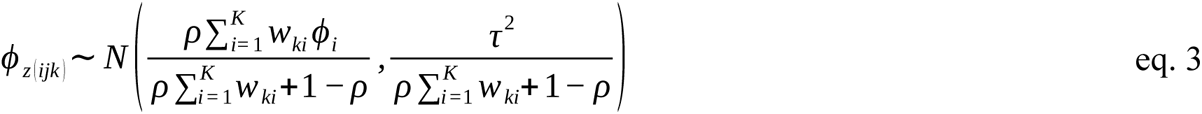

All analyses were performed using R software and the CARBayes package (Lee, 2013). Five independent chains were run; the Geweke coefficient was used to assess the convergence of the different simulations. Each chain was run for at least 3,500,000 iterations with a burnin period of 500,000 iterations and a thinning of 200 in order to avoid any correlation in the final sample. All chains were then concatenated for each parameter to form the final distributions. To provide an estimate of each parameter, the median and the 95% credible interval defined by the 2.5% and 97.5% quantiles were calculated. An effect was considered to be significant when the 95% Credible Interval (CI) did not contain zero. The median of the posterior distribution predicted by the fitted models was used to characterise each trait for each inbred line-sowing date combination.

#### Linking infestation dynamics and plant phenology

In order to investigate the effect of plant development and phenology on the dynamics of European corn borer infestation, we studied the relationship between each infestation parameter (*i_1_*, *i_2_*– *i_1_*, *r_1_*, *r_2_*, *t_0_*, *t_1_*, and *t_2_*) and the plant development and phenology traits FD and TTF. Plant development at different dates was characterized by computing the ratio between the thermal time at that date and the TTF. This ratio was computed at the beginning of the first pest generation (DEVSTAGE1) and the second pest generation (DEVSTAGE2). To estimate the start date of the first and second waves, we aggregated the raw data over the three sowing dates and over all inbred lines and chose the week where the exponential phase of the first and second generation began. Both linear and non-linear relationships were investigated with the two following models:

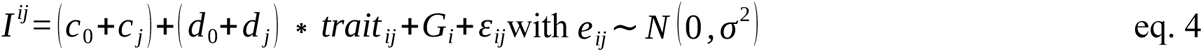

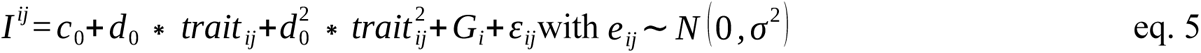

I_ij_ is the value of an infestation parameter (*i_1_*, *i_2_* – *i_1_*, *r_1_*, *r_2_*, *t_1_*, or *t_2_*) for inbred line *i* sown at sowing date *j* estimated with eq. 2; *trait_ij_*is the value of a biological trait (FD, TTF, DEVSTAGE1, DEVSTAGE2) for on inbred line *i* sown at sowing date *j* estimated with eq. 2. The parameters of these two models are: *c_0_* the intercept, *c_j_*the effect of sowing date *j* on the intercept, *d_0_* the slope of the relation, *d_j_* the effect of sowing date *j* on the slope, *d_0_^2^* the quadratic component, and *G_i_* the individual effect of inbred line *i* on the intercept modeled as a Gaussian random effect of variance *σ^2^_B_*. Traits were included in separate models to avoid highly correlated co-variables. When the biological trait considered was DEVSTAGE1 and DEVSTAGE2, we omitted the effect of sowing date on the slope and the quadratic component, as DEVSTAGE is highly dependent on the sowing date (early sown plants tend to be more developed than late sown plants). The Akaike information criterion (AIC) was used to select between the linear (eq. 4) and non-linear (eq. 5) model. When the linear model was selected, statistical tests were carried out to assess the significance of the model parameters. First, we tested the null hypothesis *d_1_* = *d_2_* = *d_3_* = 0, then c*_1_* = *c_2_* = *c_3_* = 0. When the non-linear model was selected, similar statistical tests were carried out on *d_0_* and *d_0_^2^.* Statistical tests were carried out with the lmerTest package (Kuznetsova et al., 2017) and Satterwhaite’s method for approximating degrees of freedom for the F tests (Satterthwaite, 1946).

## Results

To decipher the interplay between plant and pest phenologies, a two-year field experiment was carried out on 23 maize inbred lines sown at three different calendar dates. Both plant phenology and the natural infestation dynamics of the European corn borer were monitored and characterized by different traits. The results below describe these traits and their relationship.

### Plant phenology

For the sake of simplicity, this section describes only tasseling dates as qualitatively similar results were observed for silking dates. As a reminder, for each inbred line and sowing date, the median flowering date (FD) was used to quantify plant phenology and the thermal time to flowering (TTF) was used to quantify plant earliness.

The inbred lines in the maize diversity panel we used displayed a wide range of flowering dates (Fig. 1). For both years, the sowing date effect, the inbred line effect, and their interaction in eq. 2 were significant for both the median flowering calendar date (FD) and thermal time to flowering (TTF), i.e., the95% Bayesian Credible Interval (CI) of at least one of the sowing date, inbred line and interaction terms parameters did not contain zero (95% CI shown in Fig. S4 and S5). Spatial autocorrelation within sub-blocks was detected for both TTF and FD (ρ = 0.27) in 2022 but not in 2021. For each year, the median FD and TTF are summarized in Table S1 for each sowing. The median TTF ranged from 56.1 (95% credible interval = [54.9, 57.3]) to 89.5 thermal days [88.4, 90.6] after sowing date showing the extent of the gradient of earliness of this panel. This diversity in flowering earliness, combined with delayed sowing, resulted in an extended plant phenology across each season, with a flowering period from mid-July to early September in 2021 and from early July to mid-August in 2022 (Fig. 1).The average difference in median flowering time between two successive sowing dates was ∼ 1 to 5 days in 2021 and ∼ 8 days in 2022. In 2021, the second sowing flowered approximately at the same date as the third sowing. This can be related to the cold temperatures between the two sowing dates, which slowed down the development of plants from the second sowing. For each year, we obtained an extended plant phenology over the course of the season.

**Figure 1.**
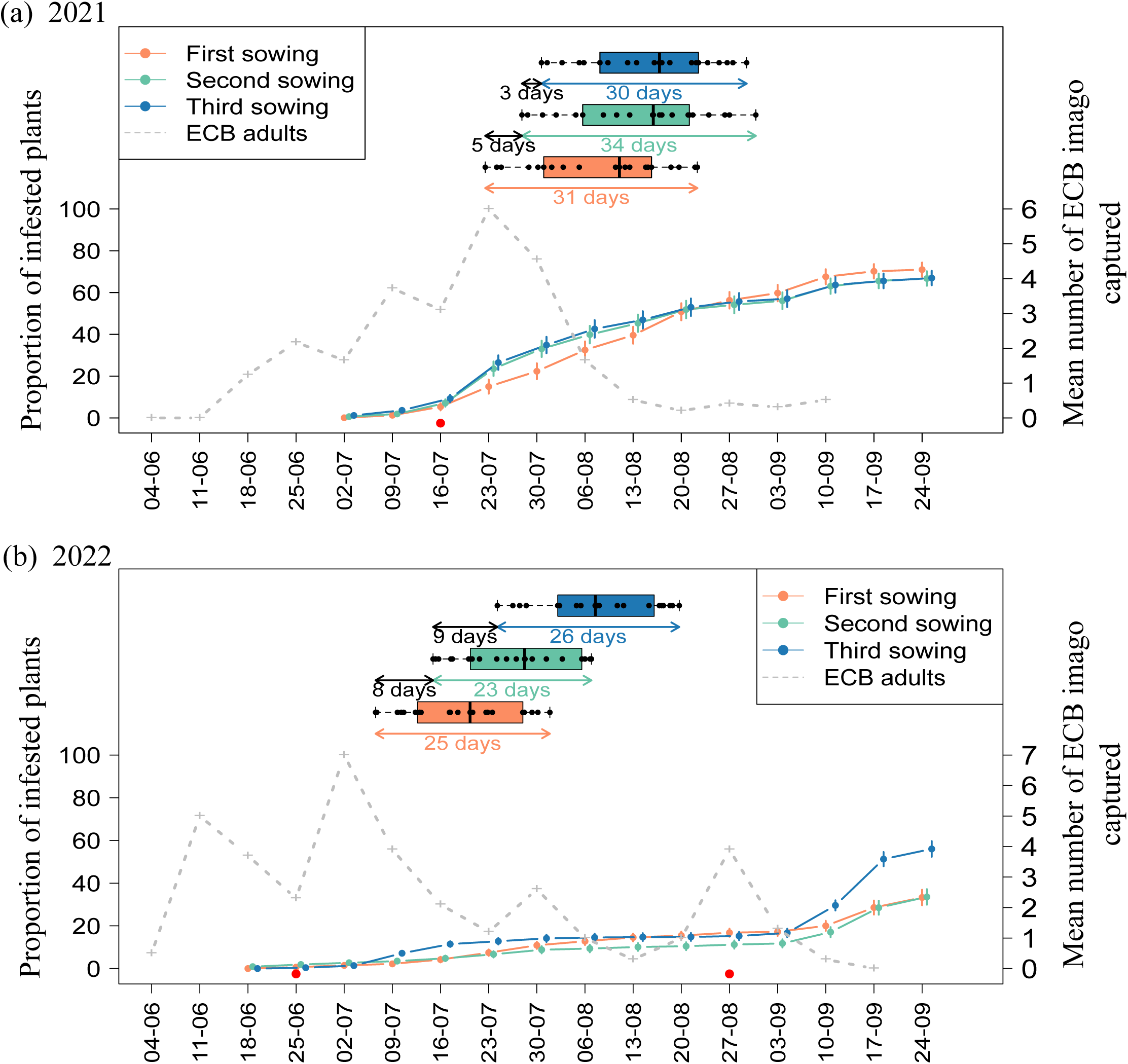
European corn borer infestation dynamics and maize phenology (FD) in 2021 (a) and 2022 (b) in Gif-sur-Yvette, Ile-de-France. The x-axis represents the week of observation. The left y-axis represents the proportion of infested plants and the right y-axis the number of European corn borer imagos captured. The mean number of infested plants from each sowing and their 95% confidence interval are shown in orange, green and blue corresponding to the first, second and third sowing, respectively. The mean number of European corn borer imagos captured in the Ile-de-France region (BSV n°32, 28/09/21 and 20/09/22) are shown in grey. Median male flowering date distributions for each sowing are shown in orange, green and blue corresponding to the first, second and third sowing, respectively. Each point within the boxplots represents the estimated median flowering date, estimated with equation 2, of each inbred line sown at a given date. The red dots represent the date at which the ratios of development (DEVSTAGE1 and DEVSTAGE2) were computed. Error bars represent the 95% credible interval for each estimate.

### Pest infestation timings in relation to plant development

The survey of pest incidence showed that one pest gest generation occurred in 2021 and two generations occurred in 2022 (Fig. 1). This is consistent with data collected in the Ile-de-France region by the national pest surveillance system (number of captured imagos documented in Bulletin de Santé du Végétal n°32, 28/09/21 and 20/09/22). As a reminder, for a given inbred line sowed at a given date, the traits DEVSTAGE1 and DEVSTAGE 2 quantify the average plant developmental state at the onset of the infestation wave caused by the first and the second generation of the European corn borer, respectively. DEVSTAGE 1 and DEVSTAGE 2 assume values below 100% when the wave started before flowering and over 100% when it started after flowering. In 2021, the infestation by the single pest generation started before maize flowering, continued throughout maize development and ended after that all maize inbred lines had flowered (Fig. 1a). The median DEVSTAGE1 was 71,5% [71.2, 71.9] for the first sowing date, 64.7% [64.4, 65.1] for the second sowing date, 61.2% [60.8, 61.5] for the third sowing date. In 2022, the infestation by the first pest generation started before maize flowering and ended when most inbred lines were still in vegetative stage, except for some early inbred lines that had already reached the flowering stage (Fig. 1b). The median DEVSTAGE1 was 52.9% [52.4, 53.4] for the first sowing date, 40.4% [40.1, 40.7] for the second sowing date, 25.3% [25.1, 25.3] for the third sowing date. The infestation by the second pest generation started when all inbred lines had flowered and lasted during maize maturation phase. The median DEVSTAGE2 was 160.8% [159.3, 162.3] for the first sowing date, 144.5% [143.4, 145.7] for the second sowing date, 128.4% [127.6, 129.3] for the third sowing date. Thus, we observed two contrasting timings of pest infestation with respect to inbred lines development between the two generations of European corn borer in 2022.

### A contrasted susceptibility between inbred lines

The median pest incidence by inbred line for each sowing was used to quantify inbred lines susceptibility. In 2021, we quantified the final incidence of the single pest generation (*i_1_*) (eq. 1). In 2022, we quantified the incidence of the first pest generation (*i_1_*) and of the second pest generation (*i_2_* -*i_1_*) (eq. 1). In both years, the inbred line effect and its interaction with sowing date were significant with respect to the pest infestation incidence of each generation ((*i_1_*) in both years, (*i_2_* – *i_1_*) in 2022, Fig. S6 and S7). Depending on inbred line and sowing dates, the median final incidence of the first generation (*i_1_*) ranged from 36% to 91% in 2021 (Fig. 2a) and 2% to 41% in 2022 (Fig. 2b). The incidence due to the second generation (*i_2_* – *i_1_*) ranged from 2% to 77% in 2022 (Fig. 2c). Some of the inbred lines of the panel were previously characterized in the literature for their cell wall components (Cm484, F271, F66), their tolerance (F618) or their susceptibility (Mo17, B73). Cm484, F271, F66 were one of the least infested in both seasons (Fig. 2). Although defined as tolerant, F618 was among the least infested lines for the first pest generation in 2022 (Fig. 2b) but not for the two other documented infestation waves (Fig. 2a and Fig. 2c). The susceptibility of Mo17 and B73 varied between years and the sowing date (Fig. 2). In 2021, Mo17 and B73 were not among the most infested lines (Fig. 2a). In 2022, they were among the least infested by the first pest generation (Fig. 2b) but were the most infested by the pest second generation, particularly in the third sowing (Fig. 2c). The variability in inbred lines susceptibility highlights the genetic diversity of the maize panel. To explain further the significant interaction effect between the inbred line and the sowing date on infestation incidence, we explored the infestation dynamics patterns from which these incidences result.

**Figure 2.**
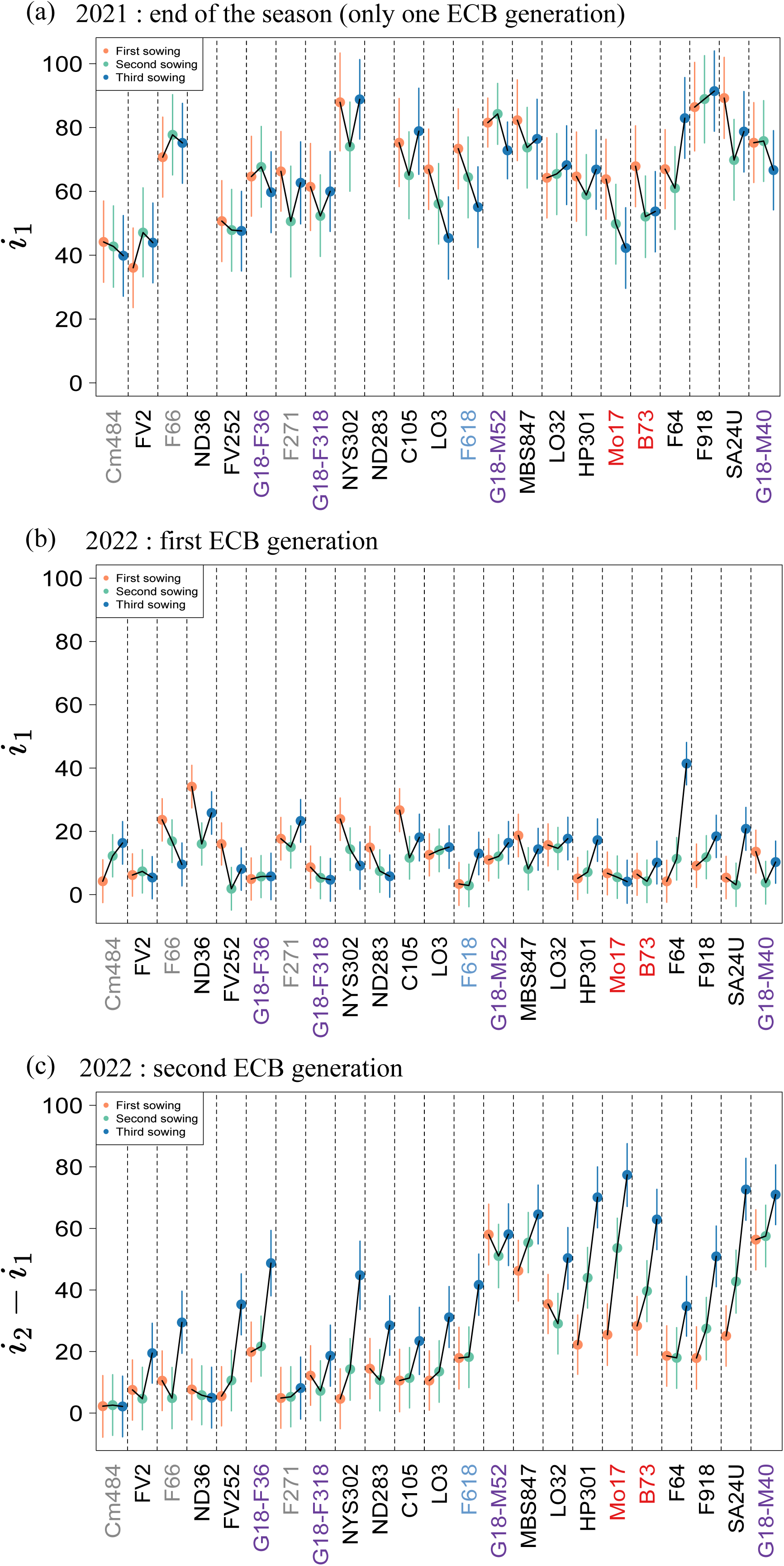
Median proportion of infested maize plants by European corn borer by inbred line and sowing date in 2021 and 2022 in Gif-sur-Yvette, Ile-de-France. Inbred lines are ordered by earliness (from early to late) as observed in the field in plants from the first sowing in 2021. Each point represents the estimated median proportion of infested plants for each inbred line sown at different dates (for both years, the first, second and third sowings are in orange, green and blue, respectively). Inbred lines in grey (Cm484, F66, and F271) have been studied for their cell wall digestibility (El Hage et al., 2018). Inbred lines in purple (G18-F36, G18-F318, G18-M52, and G18-M40) are issued from a divergent selection experiment on flowering time (Durand et al., 2010). The inbred line in blue (F618) is known for its tolerance to European corn borer (Panouillé et al. 1998). Inbred lines red (Mo17, B37) are known for their susceptibility to European corn borer (Willmot et al. 2004). (a) Proportion of plants infested by the single pest generation *i_1_* in 2021. (b) Proportion of plants infested by the first generation of borers *i_2_* in 2022. (c) Proportion of plants infested by the second generation of borers *i_2_*-*i_1_*in 2022. Error bars represent the 95% credible interval for each estimate.

### Infestation dynamics patterns

For each plot, the infestation dynamics were characterized with eq. 1 using several parameters summarizing the timing and intensity of the infestation by each pest generation. As a reminder, *t_0_* is the date of first attack on the plot, *t_1_* and *t_2_* represent the inflection points of the incidence during the first and second waves, respectively; *r_1_* and *r_2_* represent the slopes of the first and second waves, respectively. The quantities *i_1_* and *i_2_ – i_1_* correspond to the maximum incidence due to the first and second pest generation, respectively. The median of each parameter was then estimated with the model described by eq. 2 to characterize infestation dynamics for each inbred line for each sowing date.

In 2021, one single infestation wave spread over the season was observed corresponding to a single pest generation. Different infestation dynamics patterns were observed between sowing dates. On average over all inbred lines, both *t_0_*, *t_1_* and *r_1_* were significantly higher in the first sowing meaning that this sowing was infested later in the season; however incidence increased more rapidly in the first sowing than in later sowing dates (95% CI shown in Table 2 and Fig. S8). The final incidence of this single pest generation, *i_1_* was slightly higher in the first sowing than in the two others (see Fig. 1a, Table 2 and Fig. S8). Also, infestation dynamics varied between inbred lines: the inbred line effect was significant for the incidence *i_1_* (as stated in the precedent subsection), the inflection date *t_1_* and the infestation growth rate *r_1_*, and the interaction between inbred line and sowing date was significant for *i_1_* and *t_1_* (see Fig. S6 and S7). This interaction was particularly noticeable for *i_1_,* indeed, although a positive additive effect of the first sowing has been estimated, the most infested sowing was the first only for 47% of lines, the second for 24%, and the third for 29%.

**Table 2.** Median and 95% high density interval of each of the infestation parameters of European corn borer on maize by sowing date for each year (2021,2022) in Gif-sur-Yvette, Ile-de-France.

| Year | Infestation parameter | 1<sup>st</sup> sowing date | 2<sup>nd</sup> sowing date | 3<sup>rd</sup> sowing date |
| --- | --- | --- | --- | --- |
| 2021 | t<sub>0</sub> | 28.8 days<br>[27.2,30.5] | 23.7 days<br>[22.1,25.3] | 19.5 days [17.9,21] |
| 2021 | t<sub>1</sub> | 47.7 days<br>[45.7,49.8] | 42.6 days [40.6, 44.7] | 40.9 days<br>[38.9,42.9] |
| 2021 | r<sub>1</sub> | 3.0 [2.0, 4.0] | 2.6 [1.6, 3.6] | 1.2 [0.23, 2.8] |
| 2021 | i<sub>1</sub> | 68.5% [64.4, 72.6] | 63.1% [58.9, 67.4] | 64.6% [60.6, 68.7] |
| 2022 | t<sub>0</sub> | 23.9 days [20.3, 27.5] | 28.5 days [25.0, 32.2] | 24.1 days [20.3, 27.7] |
| 2022 | t<sub>1</sub> | 26.2 days [24.1, 28.2] | 19.3 days [17.2, 21.4] | 16.8 days [14.9, 18.7] |
| 2022 | r<sub>1</sub> | 3.1 [2.5, 3.7] | 3.0 [2.4,3.7] | 3.5 [2.9, 3.5] |
| 2022 | i<sub>1</sub> | 12.7% [11.2, 14.2] | 9.2% [7.7, 10.7] | 14.3% [12.8, 15.9] |
| 2022 | t<sub>2</sub> | 67.8 days [64.7, 71.11] | 78.3 days [75.5, 81] | 78.5 days [76.1, 80.8] |
| 2022 | r<sub>2</sub> | 2.25 [1.5, 3.5] | 3.3 [2.3, 4.2] | 3.8 [3.0, 4.5] |
| 2022 | i<sub>2</sub>-i<sub>1</sub> | 20%[16.8, 23.3] | 23.8% [20.1, 27.1] | 41.2% [38, 44.5] |

In 2022, two infestation waves corresponding to two generations of pests were observed with different infestation dynamics patterns between sowing and inbred lines. As in 2021, we observed differences between sowing dates. For the first pest generation, the infestation of the second sowing occurred significantly later (*t_0_*) and was less intense than that of the first and third sowing dates (*i_1_*) (95% CI shown in Table 2 and Fig. S9). No significant difference was found between the infestation growth rates (*r_1_*) of the different sowing dates (95% CI shown in Table 2 and Fig. S9). For the second pest generation, on average over all inbred lines, the third sowing was significantly more than twice as infested as the first two sowing dates (*i_2_* – *i_1_*), and was more rapidly infested (*r_2_*) (95% CI shown in Table 2 and Fig. S9). The median incidence *i_2_* – *i_1_* in the third sowing was 41.2% [38,44.5] compared to 20% [16.8, 23.3] and 23.8% [20.1, 27.1] in the first and second sowing. At the end of the season, the third sowing was the most infested considering the accumulation of the two waves of infestations (Fig. 1b). We were able to classify each plot in 2022 as being infected by larvae from the first generation, the second generation, or both generations by using the fit of the incidence at the plot level. The distribution of these different infestation dynamics depended on the sowing date (Chi-squared statistic= 36.95; df = 4; P < 0.001) with 74% of plots sown at the third sowing date being mainly infested by both pest generations as opposed to 40% of plots sown at the first and second sowing dates (Fig 3a). Moreover, we observed contrasting patterns between inbred lines with the inbred line effect being significant for the timing (*t_1_*, *t2*), the growth rate (*r1*, *r2*) and the incidence (*i_1_*, *i_2_*-*i_1_*) of both infestation waves (95% CI shown in Fig S9). We found that early inbred lines seemed more susceptible to first-generation larvae (first infestation wave), while late inbred lines seemed more susceptible to larvae from both generations (Fig. 3b). As in 2021, the interaction between the inbred line and the sowing date was significant for the incidence of both generations (*i_1_* and *i_1–_ i_2_*)

**Figure 3.**
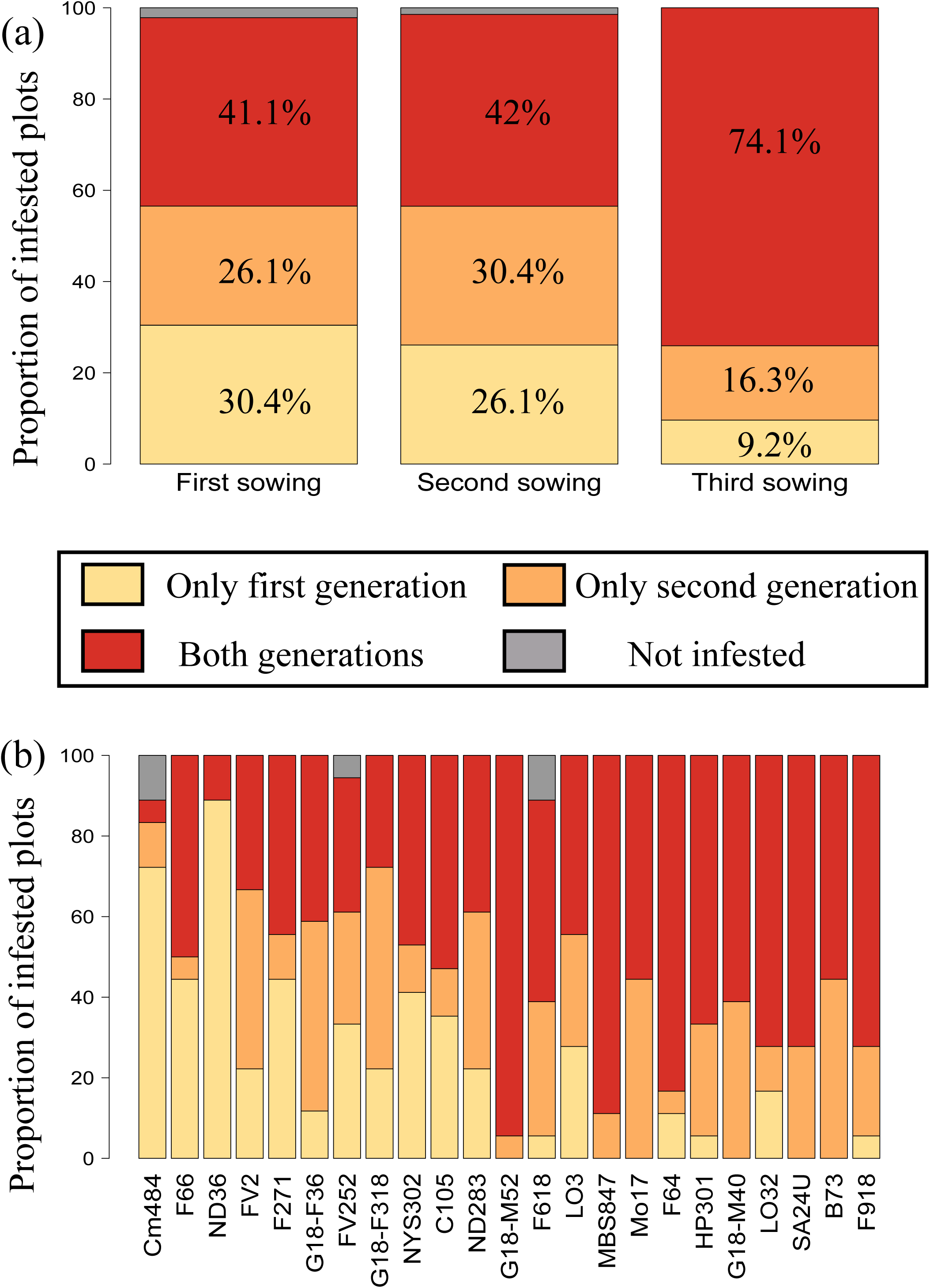
Proportion of maize plots infested by the first generation, the second generation, both generations of European corn borer and not infested in 2022 in Gif-sur-Yvette, Ile-de-France. The proportion of plots in each category is represented by sowing date in (a), and by inbred line in (b). Inbred lines are ordered by precocity (from early to late) as observed in the field in plants from the first sowing.

Overall, we observed an effect of both the sowing date, the inbred line and their interaction on the dynamics of pest attack for each wave of infestation and each year. The susceptibility of an inbred varied depending on the sowing date and the wave of infestation suggesting an effect of plant development. To explain this pattern further, we explore below whether the incidence after the first and second generation can be explained by plant phenology and development at the onset of the attack.

### Relationship between plant development and phenology and infestation dynamics

In order to determine whether infestation dynamics is impacted by a match-mismatch between plant and pest phenologies, we studied the relationship between infestation parameters and plant phenology, plant earliness and plant development at the onset of each infestation wave As a reminder, we characterized each inbred line sowed at a each date by its phenology (FD), its earliness (TTF) and its development state at the start of infestation waves (DEVSTAGE1 and DEVSTAGE2).

In 2021, the timing (*t_1_*) and incidence (*i_1_*) of the pest attacks varied linearly with phenology (FD) (*t_1_* : F=51.87; df=1, 54.68, P < 0.001; *i_1_* :F = 7.29, df=1, 22.93, P =0.012), earliness (TTF) (*t_1_* : F =24.98, df =1, 22.03, P < 0.001; *i_1_*: F=6.24, df=1, 22.41, P=0.020) and developmental state (the fraction of development DEVSTAGE1) at the onset of the infestation (*t_1_* : F=11.99, df = 1, 35.59, P< 0.001; *i_1_*: F =7.11, df=1, 23.67, P=0.013) (Fig. S10 and S11, Fig. 4a, and Table S2). The most mature plants at the time of arrival of the pest were infested later and to a lesser extent than less developed plants. However, this relationship is blurred by the significant inbred line effect, suggesting differences in genetically determined resistance factors between inbred lines.

**Figure 4.**
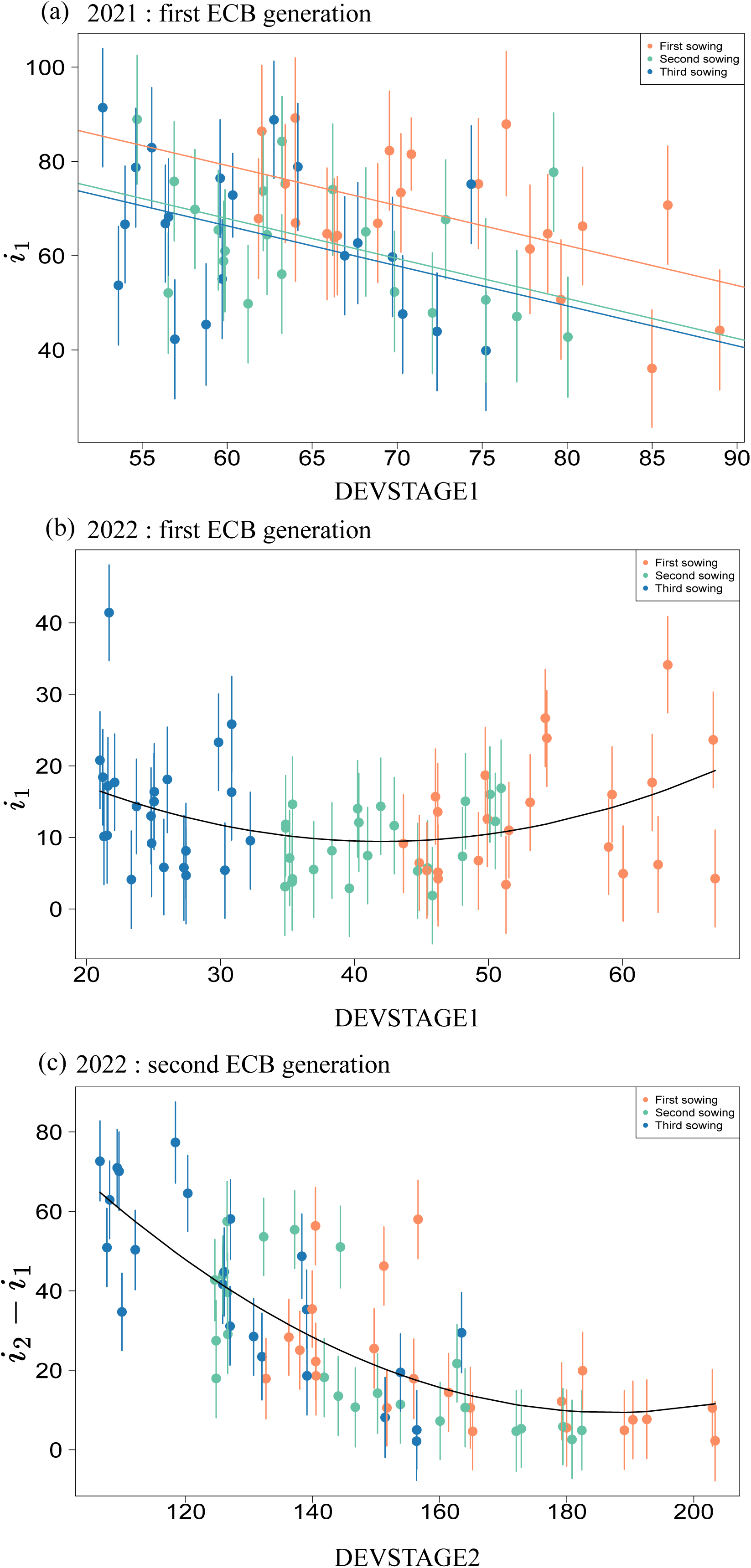
Relationship between the median proportion of infested maize plants by European corn borer and maize plant development in 2021 and 2022 in Gif-sur-Yvette, Ile-de-France. Each point represents the estimated median proportion of infested plants for each inbred line sown at different dates (for both years, the first, second and third sowings are in orange, green and blue, respectively). The x-axis represents the fraction between the thermal time to flowering of each inbred line and the thermal time associated with the date at which plant development was studied. (a) Proportion of plants infested by the single pest generation *i_1_*in relation to plant development at the onset of the 2021 infestation. (b) Proportion of plants infested by the first generation of borers *i_1_*in 2022 in relation to plant development at the onset of the first infestation wave. (c) The proportion of plants infested by the second generation of borers *i_2_* – *i_1_* in 2022 in relation to plant development at the onset of the second infestation wave. Error bars represent the 95% credible interval for each estimate.

In 2022, the level of infestation after the first pest generation *i_1_* was non-linearly related (eq. 5) to FD (Chisq = 12,47, df = 1, P<0.001) (Fig. S12a), TTF (Chisq = 4,77, df = 1, P=0.029) (Fig. S12b), and DEVSTAGE1 (Chisq = 11.89, df = 1, P<0.001) (Fig. 4b). Both the least and most developed plants were, on average, the most infested. The level of infestation after the second pest generation *i_2_* – i*_1_* was linearly related to FD (F=29.72, df=1, 23.35, P<0.001) (Fig. S13a) and TTF (F=30.02, df=1, 23.21, P<0.001) (Fig. S13b), and non-linearly related (Table S2) to DEVSTAGE2 (Chisq=30.81, df=1, P<0.001) (Fig. 4c). The linear equation (eq. 4) used for FD and TTF enabled us to take into account the effect of sowing date, which impacts FD and TTF. These results show that the most developed plants at the start of the second pest generation tended to be less infested than the least developed plants. This pattern seems better explained by the fraction of development (DEVSTAGE2) than by earliness (TTF). Indeed, when modeling the relationship between *i_2_* – *i_1_* and TTF, the strength of this relationship increased with sowing date with a correlation coefficient of 57% (t=3.12, df=21, P = 0.004) for the first sowing, and 74% (t=5.06, df=21, P <0.001) and 80% (t=6,13, df=21, P <0.001) for the second and third sowing dates, respectively. We observed that late inbred lines from the first sowing and early inbred lines from the third sowing had reached the same level of development and experienced comparable levels of infestation. These results show that the state of plant development at infestation onset affects the success of the pest, but the non-linearity of this relationship highlights the importance of the match-mismatch between plant and pest phenologies.

## Discussion

We set up a field experiment with staggered sowing dates to investigate the drivers of pest success on a panel of maize inbred lines that differ in earliness. The objective was to understand how the dynamics of plant development and plant-pest phenological match/mismatch could impact the dynamics of pest infestation. Most field experiment studies on maize have quantified pest resistance by measuring borer-induced damage at the end of the season without monitoring the infestation dynamics over the season (Schulz et al., 1997; Malvar et al., 2002; Blandino et al., 2008; Beres and Gorski, 2012; Ordas et al., 2013). The results of some of these studies suggest planting maize early and/or using resistant varieties (Schulz et al., 1997; Malvar et al., 2002; Blandino et al., 2008) while others show contradictory results on the effect of maize earliness (Beres and Gorski, 2012; Ordas et al., 2013). Our results highlight the fact that pest success can be affected by the dynamics of maize development and plant-pest synchrony. Indeed, maize defenses can vary over time and consequently the host plant that is most favorable to the pest may change during the season. Thus, monitoring infestation dynamics over the season contributes to understanding the components that determine pest success. In this study, a panel of maize inbred lines at different stages of plant development was exposed to natural infestation of pests. Infestation dynamics parameters were estimated and related to parameters characterizing plant development, earliness and phenology. With a view to mitigating pest infestations, we discuss below the various possible strategies proposed in the literature in the light of previous findings and the results of this current study. We first compare our results on the different inbred lines known to be resistant or susceptible to corn borers to previous findings. We then discuss the impact of the sowing date on the pest success for the different infestation waves. We also comment on the link between earliness and plant resistance to corn borers. Finally, we discuss how the contrasting results we obtained on the effect of the inbred lines and the sowing date can be related to both maize earliness and development state at the onset of pest attacks.

Regarding the use of resistant varieties in the literature, pest tolerance, resistance, or cell wall components of some of the inbred lines included in the panel have been previously studied using different approaches. Notably, inbred line F618 was found to be resistant, with low damage scores at maturity (Panouillé et al., 1998) and low leaf palatability (Sanane et al. 2023) and the three inbred lines Cm484, F66, and F271 are known to have stems with a high lignin content (El Hage et al., 2018), which is a known defense trait. In contrast, B73 and Mo17 are known to be susceptible to European corn borer attacks, as shown by the length of the tunnels made by borers in several environments (Willmot et al., 2004). Mo17 displayed a variable level of resistance to tunneling depending on the environment, whereas B73 was among the most susceptible genotypes in all environments. The results of our study are partly consistent with these previous findings on resistant and susceptible inbred lines. Indeed, regarding inbred lines documented as resistant lines, Cm484, F66, and F271 were one of the least infested in both 2021 and 2022. F618 was one of the least infested lines by the first generation in 2022 but this finding was not repeatable over the two other infestation waves. Regarding inbred lines documented as susceptible lines, we also observed contrasting results depending on the year and the pest generation. In 2021, Mo17 and B73 were not among the most infested lines, and in 2022, they were among the least infested by the first pest generation but were the most infested by the second generation. Hence, our study provides new data on the plasticity of the inbred lines in this panel concerning their susceptibility to European corn borer attack. These discrepancies between years and previous studies may be due to various reasons, including the nature of the trait used to quantify resistance and the panel of lines to which these lines are compared, but also to the pest population and the plant/pest phenology match/mismatch.

Regarding the impact of the sowing date on pest infestation mitigation, we indeed found that infestation dynamics depended on the sowing date. For infestation by uni-voltine European corn borer, early sowing dates were found to be more susceptible (Balachowsky, 1966). In this present study, in 2021, where only one pest generation was observed, the first sowing was the most infested, in line with this previous study. Little difference was found between the two other sowing dates, which could be explained by the fact that plants from these two sowing dates had similar developmental states. The peak of the first generation in 2021 occurred when most of the inbred lines were in a vegetative stage; developmental differences state may have been irrelevant to European corn borer preferences. In 2022, two successive infestation waves were observed. In experimental studies, first generation damage is seldom recorded as most studies measure borer-induced damage at the end of the season. One exception is the study by Balachowsky (1966), which described the effect of the sowing date on European corn borer infestation notably for both bivoltine seasons. For the first pest generation, April sowing dates are described as being more susceptible whereas late May sowing dates are described as being less infested than early May sowing dates (Balachowsky, 1966). Such results are only partially consistent with the experiment described here. In 2022, the April sowing (first sowing) was indeed more susceptible to the first pest generation than the second sowing (early May) but the third sowing (late May) was more infested than the second. For the second pest generation (second infestation wave in 2022), our results are congruent with previous studies showing that second generation of European corn borer is more abundant in late sowings (Balachowsky, 1966, Malvar et al., 2002; Blandino et al., 2008) and prefer to lay their eggs and attack late sowings (Beard, 1943; Spangler and Calvin, 2000).

Regarding the link between earliness and plant resistance to corn borers, previous studies have studied the link between earliness and various resistance traits (Ordas et al., 2013, Sanane et al., 2023). Tunnel length and ear damage were shown to be less important in early flowering inbred lines than in late flowering ones (Ordas et al. 2013). Such results are also in line with results on the leaf palatability for European corn borers L2 larvae measured in (Sanane et al. 2023). Indeed, using the same collection of maize inbred lines as the one studied here, they found that, for the same development stage, late flowering inbred lines were more consumed and more rapidly consumed than early lines. This is also in line with analyses of the maize genome which have shown that QTLs for resistance to European corn borer were co-localized with QTLs for days to flowering (Jiménez-Galindo et al., 2019), although the genetic correlation between both characters was low. A significant positive genetic correlation between days to flowering and tunnel length was found in flint material (Ordas et al., 2010) and in a flint x dent population (Samayoa et al., 2014). Our results partly support a link between earliness and susceptibility to European corn borer infestation, although this link is not straightforward and depends on pest phenology. Indeed, in 2021, the correlation between final incidence and earliness was significant only for the first sowing and was weaker for the two later sowing dates. In the first sowing, early inbred lines were less infested than late lines. One explanation could be that the adult oviposition peak occurred when all inbred lines of the second and third sowing dates were in the vegetative stage, with comparable defense levels, while for the first sowing, a fraction of the inbred lines were still in the vegetative stage and others had flowered leading to contrasted levels of defense in these plants. In 2022, the relationship between the incidence of the first pest generation and earliness was not linear, and convexity was observed. Incidence was maximal in the earliest and latest inbred lines suggesting that the earliness is not the only factor impacting pest success. A similar relationship was observed between incidence and the fraction of development, with the least and most developed inbred lines being the most infested. This could be explained by the non-linearity of defense acquisition in the early stages of development (Cambier et al., 2000). The first pest generation feeds mainly on leaves and growing tassels. In early stages of development, the main defense of maize is DIMBOA, a chemical component whose concentration in leaves decreases with development (Meihls et al., 2012). At the onset of the infestation, the least developed lines may not produce other complementary defenses, while the most developed lines might not produce DIMBOA anymore. Inbred lines at an intermediate stage of development may combine DIMBOA with other complementary mechanisms, which may explain why they were less infested, and underlines the role of a phenological match between plant and pest in the success of the pest. For the second pest generation in 2022, the correlation between earliness and incidence was significant for each sowing, with early inbred lines being less infested than late inbred lines. However, the correlation was weak for the first sowing and stronger for the second and third sowing dates. All inbred lines had already flowered and were no longer in the vegetative stage when the second pest generation arrived in the field. Cell wall composition changes with development, particularly lignin composition, which limits pest success (Sanane et al. 2023). In the first sowing, late inbred lines were sufficiently lignified to prevent young larvae from feeding and not suffer from pest infestation; indeed they showed similar infestation levels as early lines. This explains why there is only a weak relationship between the incidence of the second pest generation and earliness in the first sowing. In the delayed (second and third) sowing dates, late inbred lines were less developed and thus could be preferred by the pest compared to early inbred lines. This could explain the higher correlation between incidence and earliness in later sowing dates.

In this study, we have highlighted a variability in the dynamics of the pest infestation, which is influenced by plant development. The early flowering lines, developing faster than the late flowering lines, were less infested at the end of the season. However, pest incidence in late lines can be limited by sowing them early so that they are at an advanced stage of development when the pest arrives. Nevertheless, the relationship between plant development and pest incidence depended on the timing of pest infestation, as a nonlinear relationship was observed when the infestation by the first generation of borers occurred early in the season when all plants were in the vegetative stage. These results highlight the importance of studying plant-pest phenological match/mismatch to propose relevant control strategies.

## Supporting information

All supporting tables and figures

## Acknowledgments

The authors thank all those who have contributed to setting up the experiments, data monitoring data and formatting data. The authors declare no conflicts of interest. This research was part of the PHENOFORE project financed by SEMAE. C.D, J.L, N.G, and S.R conducted the experiment. S.R analysed the data. S.R wrote the manuscript. A.B, C.D, F.R, J.L, R.A.M, conceived and designed the research. All authors contributed to the article and approved the submitted version

