## Supplementary material for "Effects of Maize Development and Phenology on the Field Infestation Dynamics of the European Corn Borer (Lepidoptera: Crambidae)": All supporting tables and figures

### Supplementary Information

#### Contents


**1 Supplementary Figure 1** **2**

**2 Supplementary Figure 2** **3**

**3 Supplementary Figure 3** **4**

**4 Supplementary Figure 4** **5**

**5 Supplementary Figure 5** **6**

**6 Supplementary Figure 6** **7**

**7 Supplementary Figure 7** **8**

**8 Supplementary Figure 8** **9**

**9 Supplementary Figure 9** **10**

**10 Supplementary Figure 10** **11**

**11 Supplementary Figure 11** **12**

**12 Supplementary Figure 12** **13**

**13 Supplementary Figure 13** **14**

**14 Supplementary Table 1** **15**

**15 Supplementary Table 2** **16**

### 1 Supplementary Figure 1

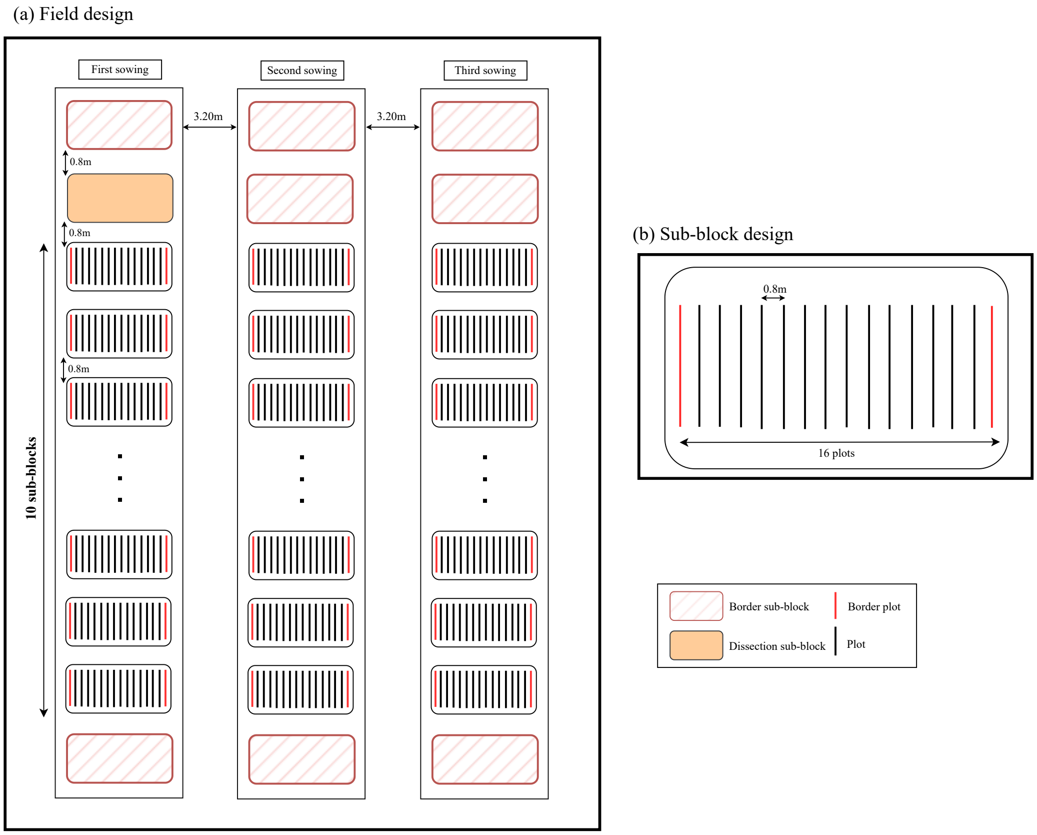


Figure S1: Design of the field experiment. (a) Global field design. Each block was associated to a sowing date and was divided into thirteen sub-blocks (0.80m apart) of sixteen plots (0.80m apart). For each sowing date, each inbred line was sown in six plots (replicates). The six replicates, for each inbred line and each sowing date, were randomized in the blocks with the condition that the same inbred line must not be sown more than twice in a sub-block. The first and last sub-block of each block were used as borders.(b) Sub-block design. Each inbred was assigned to a different plot consisting in a single row. Twenty-five seeds were sown in each plot. The first and last plot of each sub-block were used as borders.

#### 2 Supplementary Figure 2

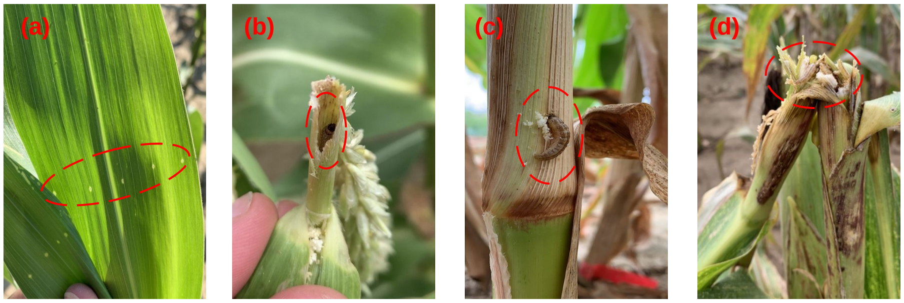


Figure S2: *O. nubilalis* damages on maize plants. (a) Holes in young maize leaves made by *O. nubilalis* larvae. (b) *O. nubilalis* larva boring into a broken panicle. (c) Frass and *O. nubilalis* larval boring into a maize stem. (d) *O. nubilalis* larva feeding in a broken maize stem.

### 3 Supplementary Figure 3

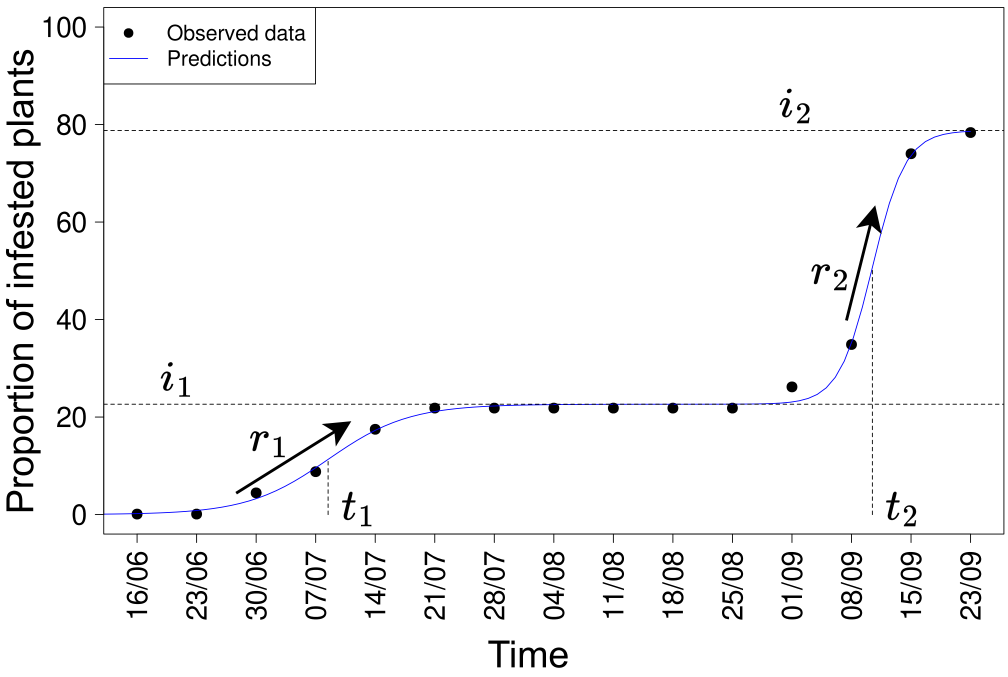


Figure S3: Double logistic curve characterization of infestation dynamics of *O. nubilalis* on a maize plot. This is an example of one infestation dynamic over a plot in 2022 by the the two successive generations of borers. The x-axis represents the observation time, the y-axis represents the proportion of infested plants. The black dots are the observed proportions of attacked plants in a plot and the blue line is predictions of the logistic growth model. $i_1$and $i_2$ are the asymptote of the first and second logistic. $t_1$ and $t_2$ are the inflection points of the first and second logistic. $r_1$ and $r_2$ are the slope parameters of the first and second logistic.

### 4 Supplementary Figure 4

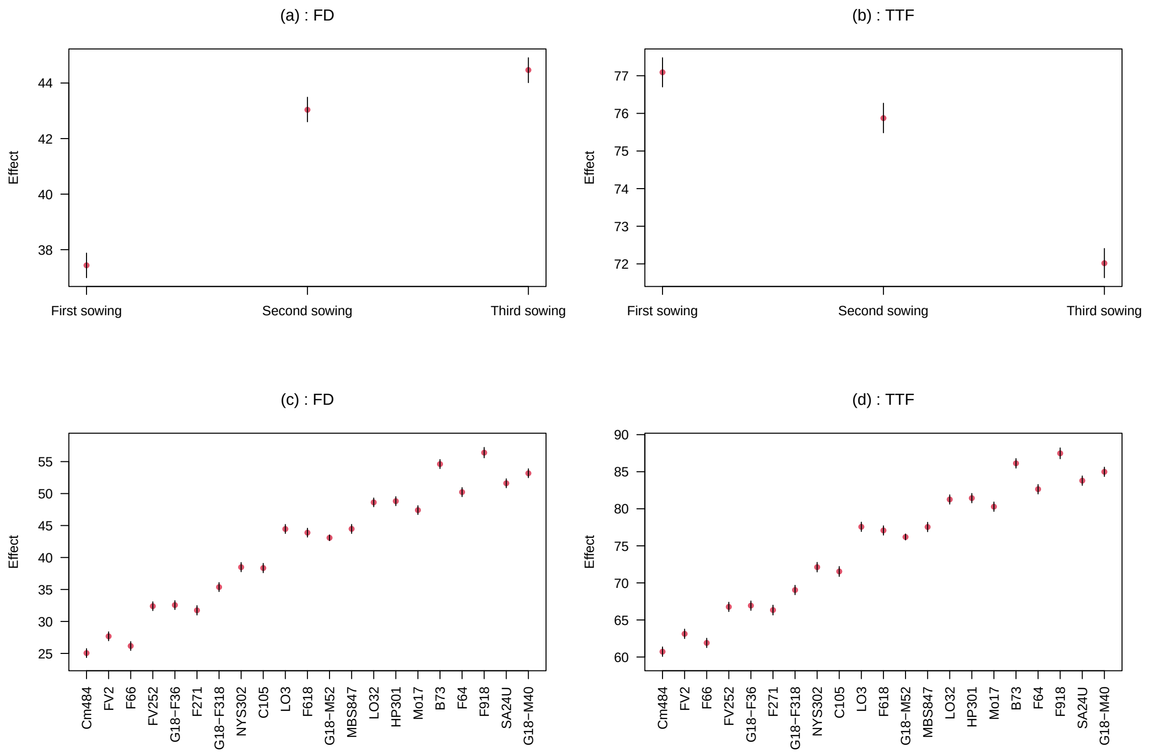


Figure S4: Estimated median of the sowing date effect $\beta$, and the genotype effect $\alpha$ for the maize flowering date (FD) and the thermal time to flowering (TTF) in 2021. Error bars represent the 95% credible interval for each estimate. Inbred lines are ordered by earliness (from early to late) as observed in the field in plants from the first sowing in 2021.

### 5 Supplementary Figure 5

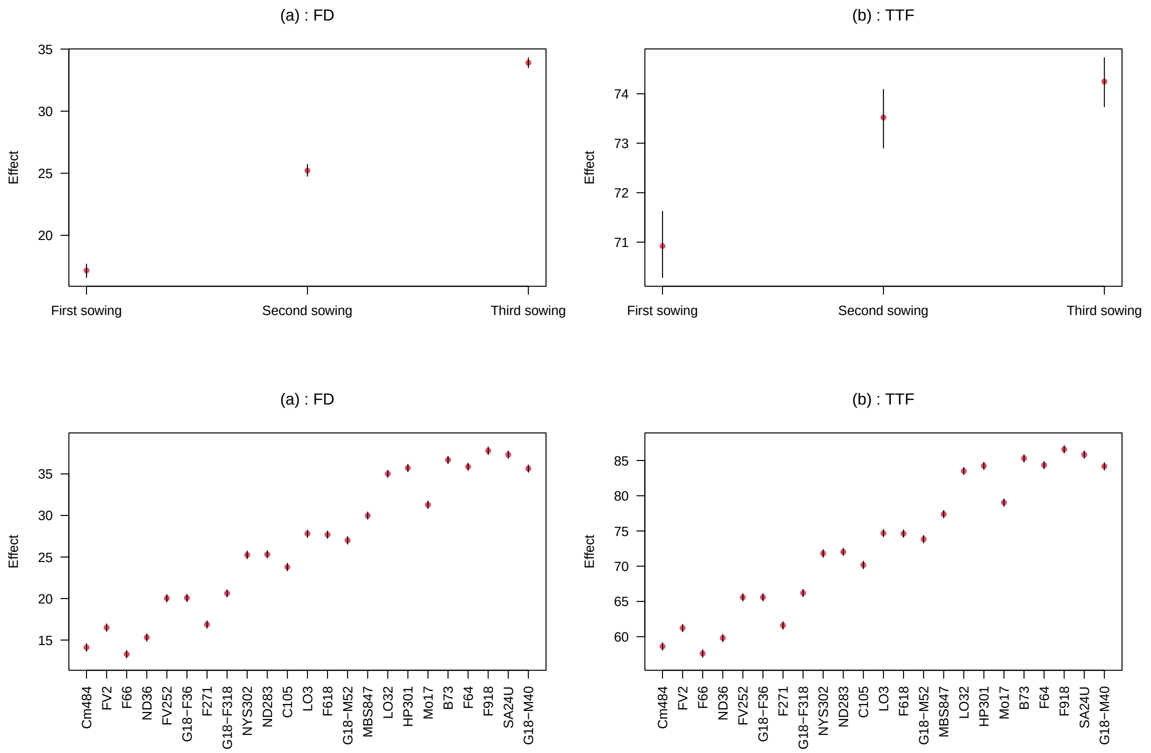


Figure S5: Estimated median of the sowing date effect $\beta$, and the genotype effect $\alpha$ for the maize flowering date (FD) and the thermal time to flowering (TTF) in 2022. Error bars represent the 95% credible interval for each estimate. Inbred lines are ordered by earliness (from early to late) as observed in the field in plants from the first sowing in 2021.

### 6 Supplementary Figure 6

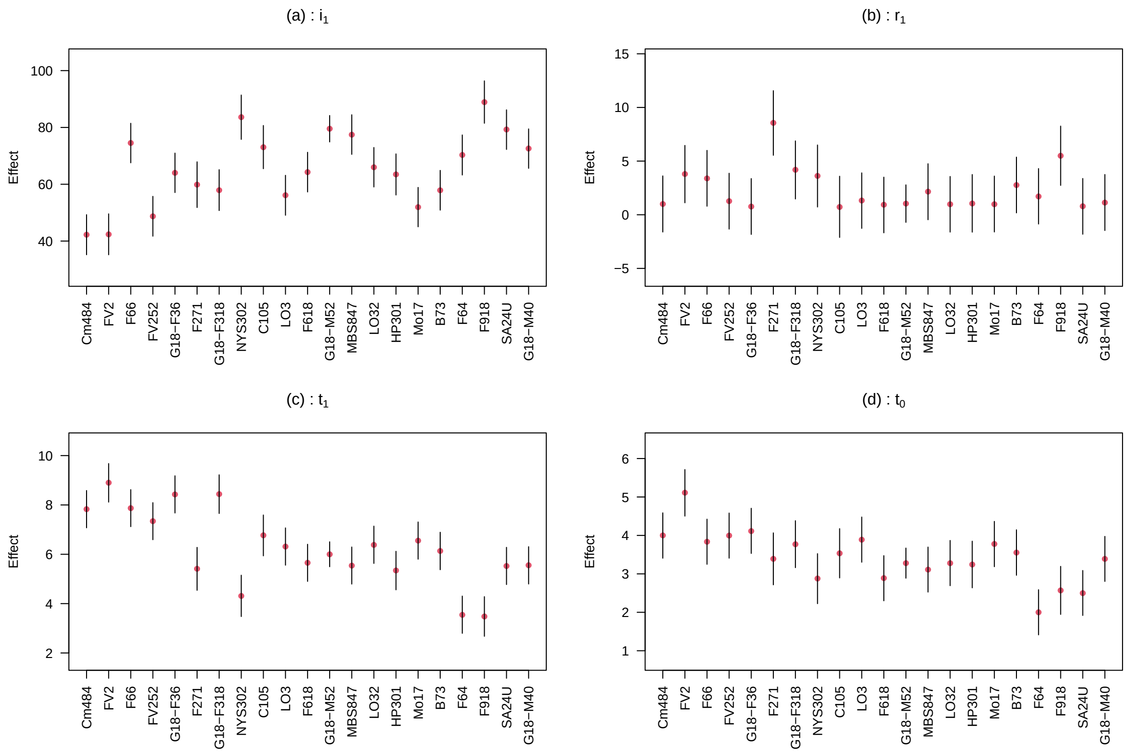


Figure S6: Estimated median of the genotype effect $\alpha$ for each infestation parameter dynamics of *O. nubilalis* in 2021. Error bars represent the 95% credible interval for each estimate. Inbred lines are ordered by earliness (from early to late) as observed in the field in plants from the first sowing in 2021.

### 7 Supplementary Figure 7

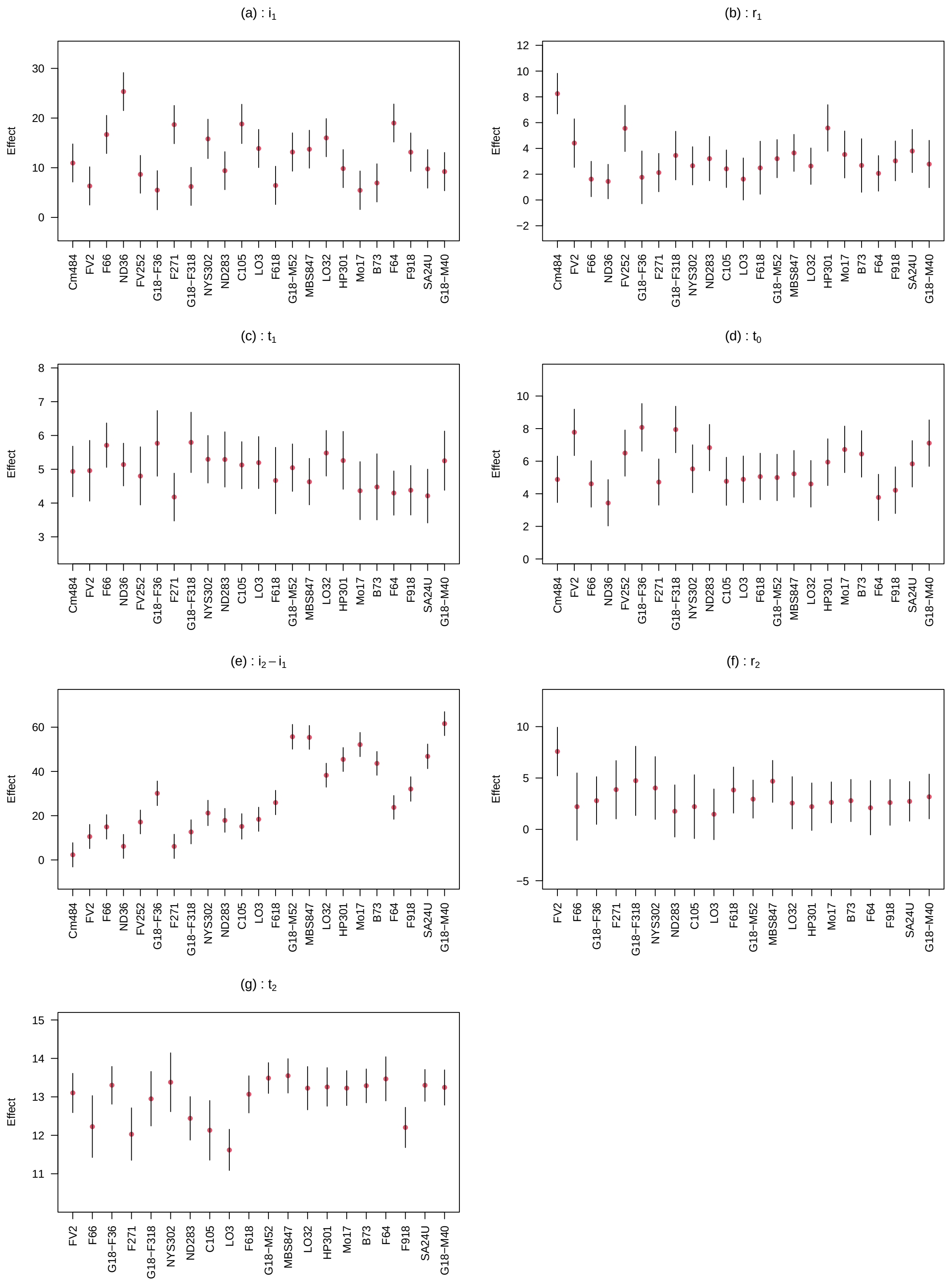


Figure S7: Estimated median of the genotype effect $\alpha$ for each infestation parameter dynamics of *O. nubilalis* in 2022. Error bars represent the 95% credible interval for each estimate. Inbred lines are ordered by earliness (from early to late) observed in the field in the first sowing date in 2021 as reference. Median for $r_2$ and $t_2$ for inbred lines Cm484, ND36, and FV252 could not be estimated as only few plots were infested by the second pest generation.

### 8 Supplementary Figure 8

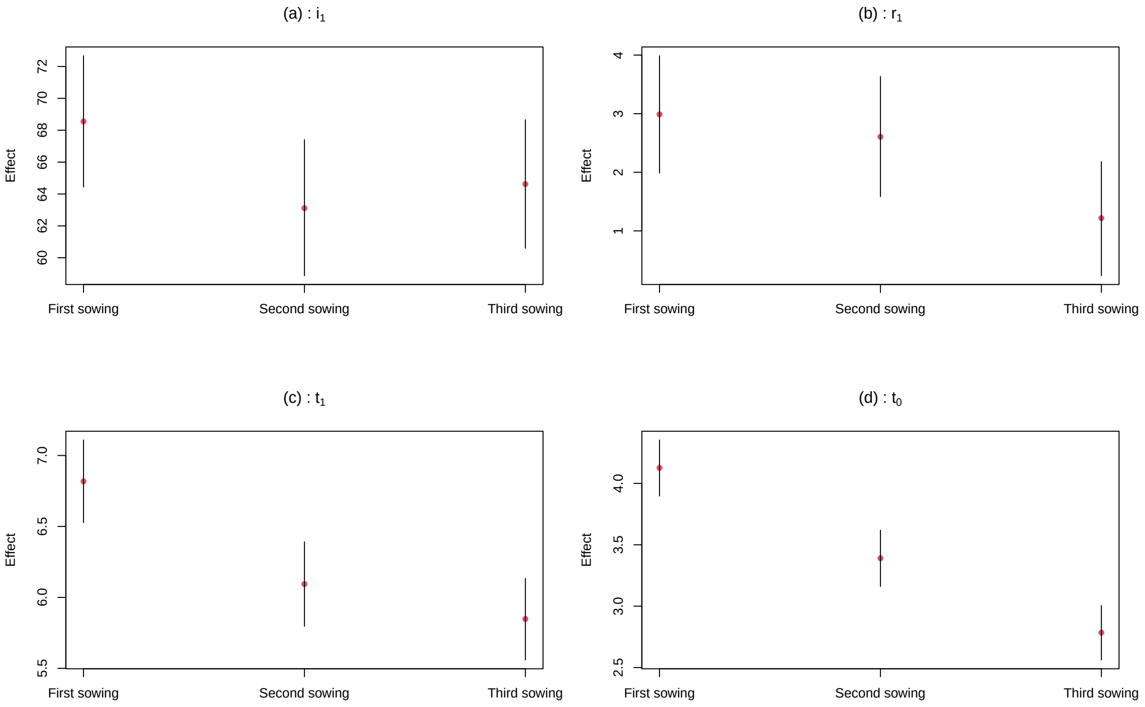


Figure S8: Estimated median of the sowing date effect $\beta$ for each infestation parameter dynamics of *O. nubilalis* in 2021. Error bars represent the 95% credible interval for each estimate.

### 9 Supplementary Figure 9

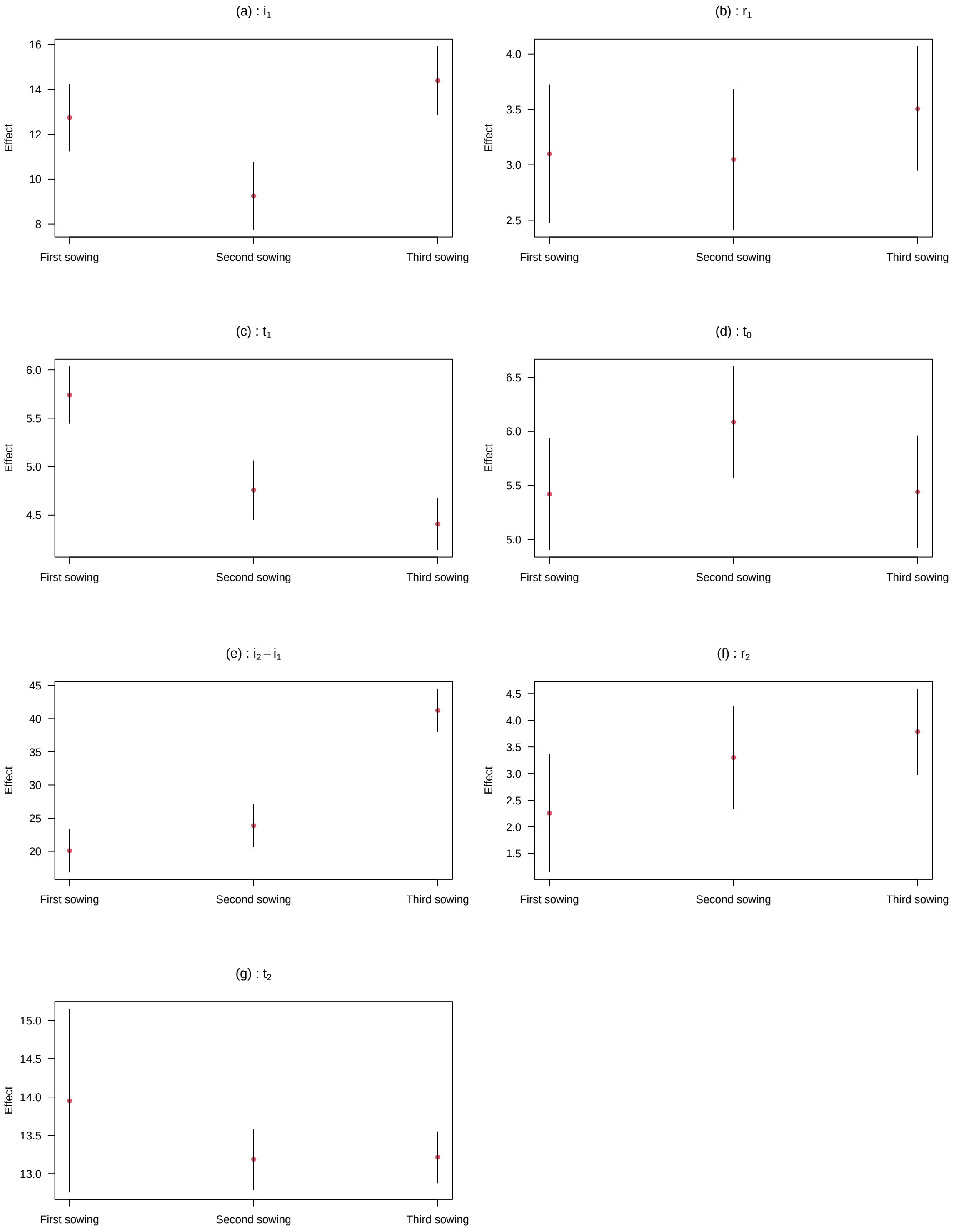


Figure S9: Estimated median of the sowing date effect $\beta$ for each infestation parameter dynamics of *O. nubilalis* in 2022. Error bars represent the 95% credible interval for each estimate.

### 10 Supplementary Figure 10

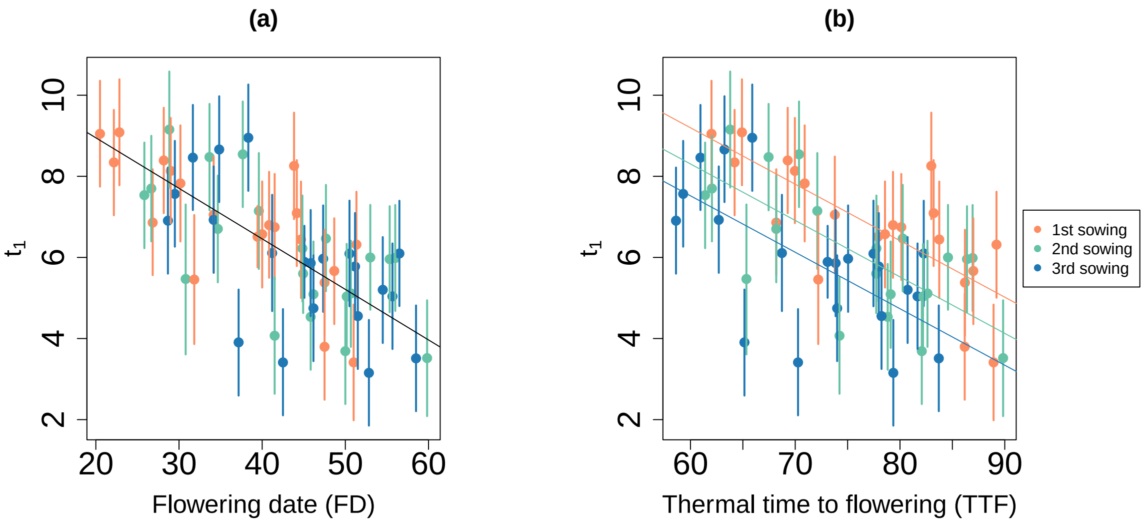


Figure S10: Relationship between the median inflection date $t_1$ of *O. nubilalis* infestation dynamics and flowering date (a) and thermal time to flowering (b) in 2021. Each point represents the estimated median inflection date for each inbred line sown at different dates (the first, second and third sowings are in orange, green and blue respectively). Error bars represent the 95% credible interval for each estimate.

### 11 Supplementary Figure 11

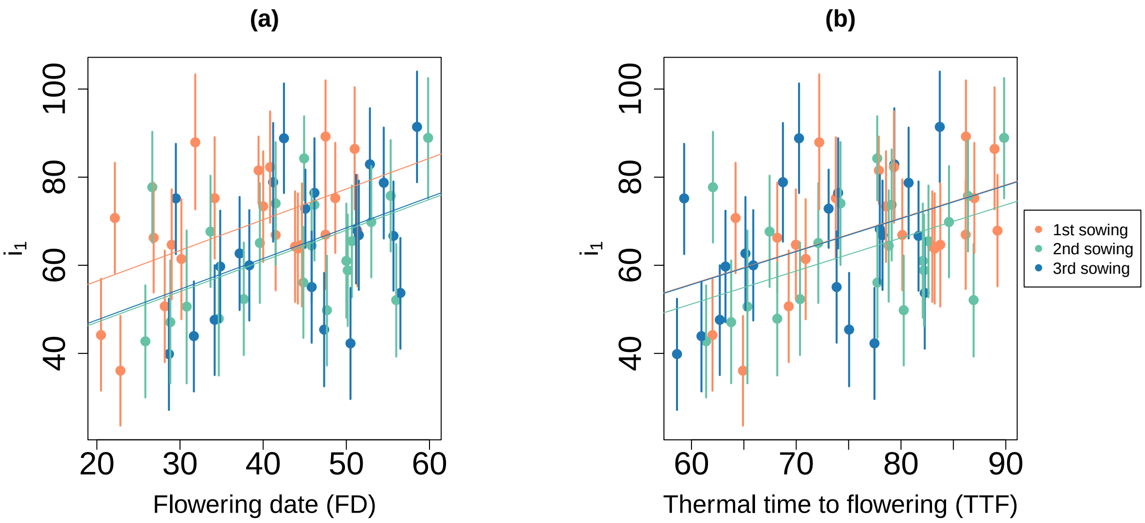


Figure S11: Relationship between the median incidence $i_1$ of the first pest generation of *O. nubilalis* and flowering date (a) and thermal time to flowering (b) in 2021. Each point represents the estimated median incidence $i_1$ for each inbred line sown at different dates (the first, second and third sowings are in orange, green and blue respectively). Error bars represent the 95% credible interval for each estimate.

#### 12 Supplementary Figure 12

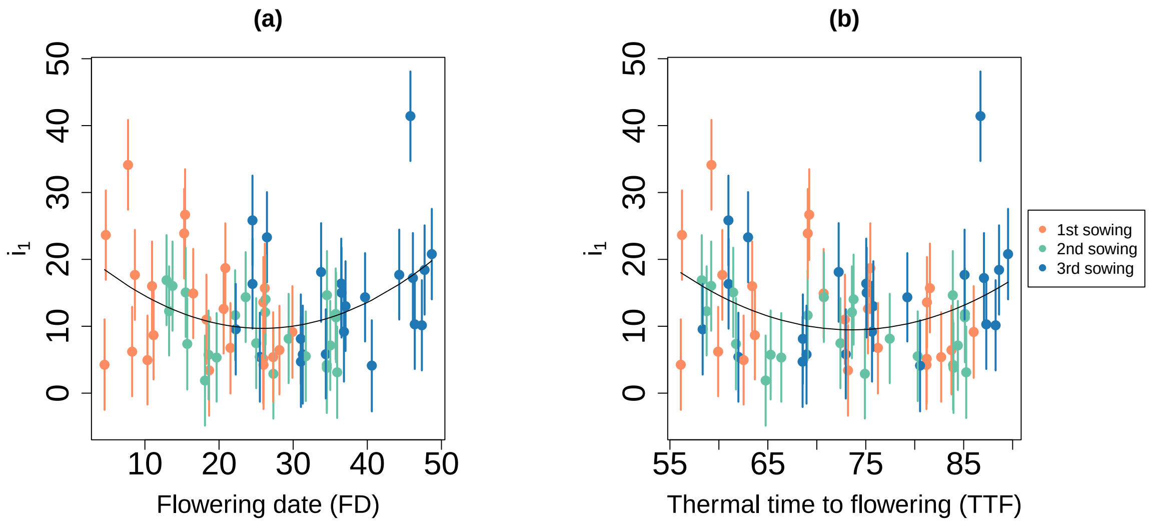


Figure S12 : Relationship between the median incidence $i_1$ of the first pest generation of *O. nubilalis* and flowering date (a) and thermal time to flowering (b) in 2022. Each point represents the estimated median incidence $i_1$ for each inbred line sown at different dates (the first, second and third sowings are in orange, green and blue respectively). Error bars represent the 95% credible interval for each estimate.

### 13 Supplementary Figure 13

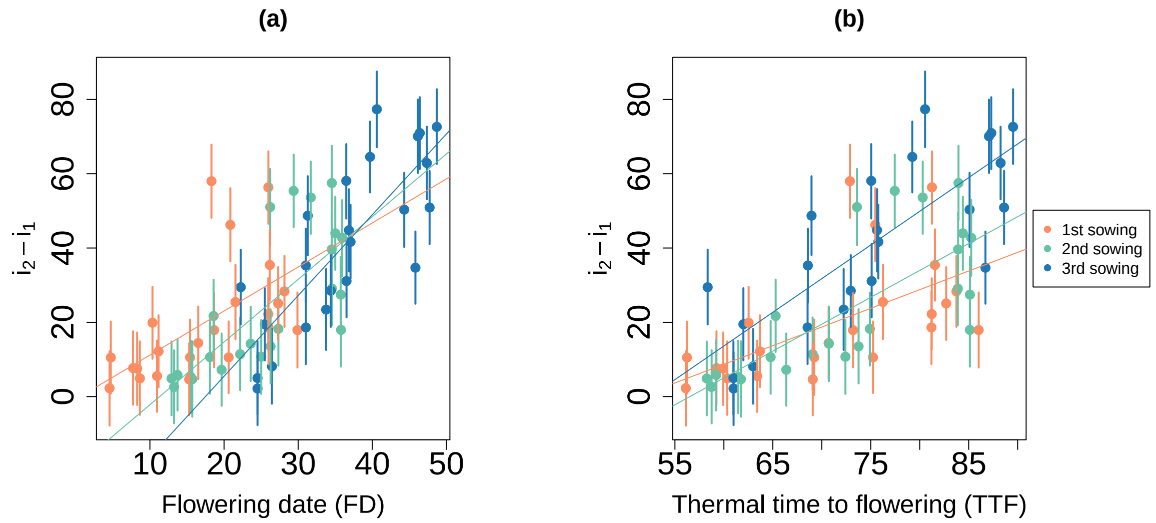


Figure S13: Relationship between the median incidence of the second pest generation $i_2$ and flowering date (a) and thermal time to flowering (b) in 2022. Each point represents the estimated median incidence $i_2$ of the second pest generation of *O. nubilalis* for each inbred line sown at different dates (the first, second and third sowings are in orange, green and blue respectively). Error bars represent the 95% credible interval for each estimate.

### 14 Supplementary Table 1

| Year | Trait | $1^{st}$ sowing date | $2^{nd}$ sowing date | $3^{rd}$ sowing date |
|---|---|---|---|---|
| 2021 | FD | $7^{th}$ August | $13^{th}$ August | $14^{th}$ August |
| 2022 | FD | $17^{th}$ July | $25^{th}$ July | $3^{rd}$ August |
| 2021 | TTF | 77.1 [76.7, 77.5] | 75.9 [75.5, 76.2] | 72.0 [71.6, 72.4] |
| 2022 | TTF | 70.9 [70.2, 71.6] | 73.5 [72.9, 74.1] | 74.2 [73.7, 74.7] |
| 2021 | DEVSTAGE1 | 71.5% [71.2, 71.9] | 64.7% [64.4, 65.1] | 61.2% [60.8, 61.5] |
| 2022 | DEVSTAGE1 | 52.9%[52.4, 53.4] | 40.4% [40.1, 40.7] | 25.3% [25.1, 25.5] |
| 2022 | DEVSTAGE2 | 160.8% [159.3, 162.3] | 144.5% [143.4, 145.7] | 128.4% [127.6, 129.3] |

Table S1 : Median and 95 % high density interval of the flowering date (FD), thermal time to flowering (TTF) and fraction of development for both years at the onset of each infestation wave (DEVSTAGE1, DEVSTAGE2) by sowing date.

### 15 Supplementary Table 2

| Year | Infestation parameter | Biological trait | $c_j \neq 0$ | $d_0 \neq 0$ | $d_j \neq 0$ | $d_0^2 \neq 0$ |
|---|---|---|---|---|---|---|
| 2021 | $t_1$ | FD | n.s | $P < 0.001$ | n.s | X |
| 2021 | $t_1$ | TTF | $P < 0.001$ | $P < 0.001$ | n.s | X |
| 2021 | $t_1$ | DEVSTAGE1 | $P < 0.001$ | $P < 0.001$ | n.s | X |
| 2021 | $i_1$ | FD | $P < 0.001$ | 0.012 | n.s | X |
| 2021 | $i_1$ | TTF | 0.044 | 0.020 | n.s | X |
| 2021 | $i_1$ | DEVSTAGE1 | $P < 0.001$ | 0.013 | n.s | X |
| 2022 | $i_1$ | FD | X | X | X | $P < 0.001$ |
| 2022 | $i_1$ | TTF | X | X | X | 0.029 |
| 2022 | $i_1$ | DEVSTAGE1 | X | X | X | $P < 0.001$ |
| 2022 | $i_2 - i_1$ | FD | $P < 0.001$ | $P < 0.001$ | $P < 0.001$ | X |
| 2022 | $i_2 - i_1$ | TTF | $P < 0.001$ | $P < 0.001$ | $P < 0.001$ | X |
| 2022 | $i_2 - i_1$ | DEVSTAGE1 | X | X | X | $P < 0.001$ |

Table S2 : Summary of statistical tests on the relationship between infestation parameters and biological traits. The values correspond to the p-values associated with the statistical tests defined by the column names. Parameters $c_j$, $d_0$, $d_j$, and $d_0^2$ correspond to the parameters of equations 5 and 6. X is written when the test is not performed. n.s is written when the statistical test is not significant.
